# In Vitro Reduction of Device Thrombosis Using a Combined Nitrite and Red Light Treatment

**DOI:** 10.1101/2023.06.10.544453

**Authors:** Elmira Alipour, James E. Jordan, Laxman Poudel, D. Clark Files, Daniel B. Kim-Shapiro

## Abstract

Device thrombosis occurs in otherwise life-saving procedures involving blood-contacting medical devices. Despite the use of systemic blood thinners, anticoagulants, and antiplatelet agents, device thrombosis can lead to substantial neurological damage, limb loss, death, and prolonged illness. Systemic treatments can also lead to bleeding. New methods to locally reduce thrombosis are urgently needed. Earlier work has shown that nitrite is a unique nitric oxide (NO) donor that is well-suited to use in blood and that its ability to inhibit platelet activation is potentiated by far-red light. In this study, we have applied our combined nitrite/light treatment in a prototypical technique used to prevent device thrombosis in extracorporeal circulation. We show that circuit pressure and survival are improved by an average of 213 percent with our treatment compared to the control. In addition, the dual therapy preserved platelet numbers at the end of the circulation time (%17 difference in platelet loss), and it reduced circuit hemolysis 2.3 fold. Thus, the combination of nitrite and red-light illumination has potential to prevent device thrombosis and to lead new clinical applications and practices.

## Introduction

Formation of blood clots is often a beneficial process to stop bleeding and repair damage after a wound. However, clot formation when there is no wound can lead to significant morbidity and mortality; this process is referred to as thrombosis. There are many types of thrombosis including deep vein thrombosis, arterial thrombosis, and device thrombosis - the focus of this study. Use of several devices including stents, pacemakers, heart valves, ventricular assist devices (VADs), central venous catheters (CVCs), dialysis circuits, cardiopulmonary bypass (CPB), and extracorporeal membrane oxygenator (ECMO) circuits for respiratory or cardiac failure as well as hemodialysis involve contact of the devices with blood which can result in thrombus formation. Device thrombosis is a significant complication in otherwise life-saving interventions.^1,2^

Such thromboses can result in thromboembolism, stroke, myocardial infarction, and device malfunction.^3^ In general, standard care has been employed to solve the problem of device thrombosis: (i) modification of the surface of the device to reduce protein adsorption, and (ii) administration of systematic anti-clotting or anti-platelet drugs. The latter solution is complicated by bleeding and, in some cases, heparin-induced thrombocytopenia or hypertriglyceridemia.^4,5,6^ However, with all of the advances made, failure rates are still as high as 6%.^2^ While the incidence of circuit clotting depends on a number of factors, one recent study found clotting in up to 38% of patients undergoing Continuous Replacement Renal Therapy (CRRT).^7^ Up to one-third of neonates and children suffer thrombosis complications from extracorporeal circulation devices.^8^ Therefore, methods to reduce circuit thrombosis would be valuable.

Substantial evidence supports the potential of nitric oxide (NO) for treatment in thrombotic conditions.^9,10,11,12,13,14^ NO reduces platelet activation,^10,12,15^ and reduces circulating blood cell adhesion to endothelia.^12,16,17,18,19^ The potential of NO to treat thrombosis is primarily through its action in reducing platelet activation,^10,20,21,22,23^ however, our recent work suggests that it will also affect other aspects of clotting.^24,25,26^ Our study is based largely on the premise that nitrite, once considered to be biologically inert in human physiology,^27^ is converted to the important signaling molecule, nitric oxide, by hemoglobin via a new function of this oxygen-carrying molecule.^28,29,30,31,32,33,34,35^ The involvement of red blood cells (RBCs) in the bioactivation of nitrite and inhibition of platelet activation through a NO-dependent mechanism has previously been shown by several investigators.^28,24,34,25,36^ Importantly, nitrite is the most suited NO therapeutic for action in blood since nitrite serves as a unique NO donor that requires red cells for bioactivation, unlike other nitric oxide donors that lose activity in the presence of RBCs.^34^

Far-red light (600-700 nm) and near-infrared light (700-1400 nm) have been used clinically for improved wound healing, tissue generation, reduction of inflammation, and increasing cerebral blood flow to improve cognition.^37,38^ Previous work has shown that far-red light has antithrombotic properties^39,40,41,42,43^ The mechanism for these actions has been attributed to increased blood flow, tissue oxygenation, and angiogenesis^44,45,46^ secondary to improved mitochondrial function attributed to cytochrome c oxidase redox state and function.^46,47,48,49,50^ However, we and others have suggested that the mechanism for beneficial effects by far-red light illumination (∼ 660 nm) relates to increases in NO bioavailability.^24,25,51^ We have shown that illumination with far-red light dramatically enhances nitrite bioactivation by red blood cells and potentiates inhibition of platelet activation, platelet adhesion, and other measures of clotting.^25,24^

We hypothesize that nitrite increases NO bioavailability through a mechanism involving RBC bioactivation, and this action is potentiated with far-red light illumination that can be used for treatments aimed at decreasing device thrombosis. A new preventative treatment using dual therapy to reduce device thrombosis has been developed and introduced in this study. We have built a prototype device and tested it against a control system with no treatment. The focus here is on device thrombosis as this type of clotting occurs at defined locations that would be accessible to illumination with red light. Device thrombosis is a complex process that involves the adhesion of blood molecules like fibrinogen and Von Willebrand factor to the device surface followed by platelet adhesion, platelet activation, platelet aggregation, and clotting.^1^ Nitric oxide is a strong anti-platelet agent, and NO production via nitrite and far-red light could provide clinical benefits in preventing device thrombosis.

## Materials and Methods

### Whole blood, Red Blood Cells, and Platelets

Whole blood, packed red blood cells, and platelet-rich plasma from healthy donors were ordered from Interstate Blood Bank, Inc (Memphis, TN) and Oneblood Company (Charlotte, NC). The units of blood components were subsequently PCR tested by IBB for HBV DNA, HIV-1 RNA, HCV RNA, WNV RNA, ZIKV RNA, and BABESIA. In the cases of negative test results, the packages were shipped overnight to the Kim-Shapiro lab in the physics department of Wake Forest University. The RBC unit was kept on ice and platelets were shipped at room temperature to preserve platelet function.

### Circuit Setup (Synchronous testing)

On the day of the experiment, two identical circuits were set up with equal lengths of tubing (Tygon, 1/4 in ID, 3/8 in OD), the same number of connection points, and similar connecting valves and injection sites. One circuit was the “untreated (control) circuit” and the other the “treatment circuit”. Both systems were connected to a Harvard Apparatus pump (PHD 2000) for intermittent infusions, the same nitrogen tank, and a Haake temperature control circulating water bath. Both circuits ran simultaneously. Figure 1 is a scheme of one of the circuits. The reservoir and the Affinity Pixie™ oxygenators were purchased from Medtronic (Minneapolis, MN). The purpose of the oxygenator was to function as a deoxygenator to bring the air-equilibrated blood products down to venous oxygen saturations. The lights are Mid panels from Mitored (Scottsdale, AZ). The oxygenator was also the primary surface to which platelets adhered and increased resistance to flow. It is the representative surface that requires local protection.

**Figure 1.**
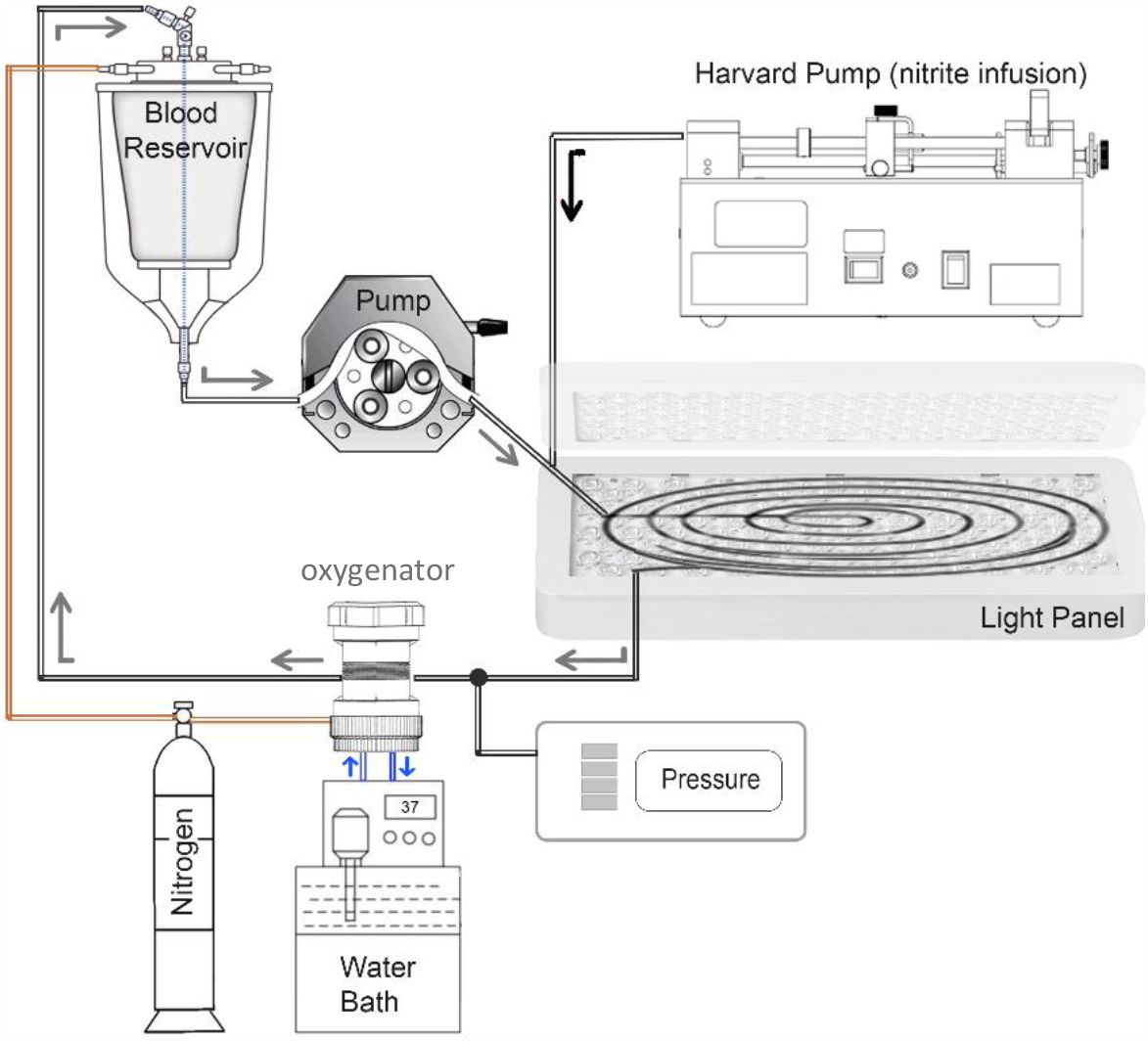
Schematic figure of our prototype. The experiment starts with the reservoir filled with 250 mL prime saline, removing oxygen using nitrogen gas flow for an hour, maintaining the temperature at 37C, and finally, 250 mL whole blood or blood mixture is added to the system. The pump leads the blood between two red light panels (sandwiched) while sodium nitrite/saline dosing is applied by a Harvard pump. The data from changes in circuit pressure is collected via a blood pressure analyzer placed before the oxygenator. While the reservoir and the oxygenator are both connected to a nitrogen gas tank, the blood goes back to the reservoir for the next cycle. The control circuit is identical except there are no light panels.

During the experiment, routine precautions were taken for blood safety, and in addition to that, the setup was originally confined and isolated by Plexiglas to minimize blood exposure. The whole setup was then moved into a biosafety cabinet for maximizing safety.

A Masterflex pump (Cole-Parmer, Vernon Hills, IL) flowed nitrogen gas through empty circuits for 20 minutes. Later, the pump was paused for the addition of 250 mL of normal saline (pH 7.4) to the reservoirs of each circuit. The pump and gas flow then continued as the circuits were maintained at 37°C temperature by the water bath. After deoxygenation (∼ 1 hour) and debubbling, an RBC-platelet mixture containing 230 mL packed RBCs, 60 mL PRP, and 230 mL saline buffer was divided into two 250 mL between reservoirs of both treated and control systems while the pump was paused. Meanwhile, the Harvard pump was also ready to inject 5 mL of 100 μM sodium nitrite and saline to the treated and control circuits respectively at the same flow rate of the blood, 150 mL/min (similar rates to that used in Continuous Replacement Renal Therapy (CRRT).

After the addition of the blood to the circuit, the first nitrite/saline infusion began immediately (zero hour) and immediately afterward both Mitored light panels started illumination in the treatment loop and remained on until the end of the experiment followed by the hourly nitrite/saline infusions. Blood samples (2 mL) from each circuit were collected hourly prior to infusing 2 mL of 100 μM nitrite/saline into the related circuits. Depending on the variations in pressure over time, the gradual addition of calcium chloride became required to initiate clotting in the systems. In those cases, calcium and/or ADP infusion started from 2.5 mM, then 0.25 mM additions every hour or every two hours depending on the rate of pressure variations. Calcium additives in the treatment circuit were mixed with the 100 μM nitrite dose. All procedures were identical for the control circuit except saline was used instead of nitrite and there was no illumination.

### Pressure Readings

Based on the chosen flow rate, it took about 3.5 minutes for one full cycle of 500 mL blood mixture. Starting from the first cycle, the pressure values (in mmHg) before and after the oxygenators were recorded every 30 minutes using a Blood Pressure Analyzer (BPA) - higher pressures indicate occlusive clotting.

### Complete Blood Count and Hemolysis Calculation

Samples (5 mL) of the first-cycled blood in both circuits were also collected in separate falcon tubes. Less than 3 mL of each was transferred into EDTA tubes for a Complete Blood Count (CBC) test performed at Atrium Health Wake Forest Baptist. To determine hemoglobin oxygen saturation, 0.3 mL of the blood samples were added to 1 mm pathlength quartz cuvettes for blood oxygenation scans on a Cary 100 spectrophotometer equipped with an integrating sphere detector to minimize light scattering effects. 1 mL was taken for comparing hemolysis using visible range spectrophotometry.

Hemolysis changes over time in the control and treated samples were determined from the supernatants in hourly collected samples. For these measurements, the blood samples were centrifuged at 300 g for five minutes and the supernatants were transferred to the 1-mm pathlength quartz cuvettes to be scanned in the visible range (450-700 nm) using a Cary 50 or Cary 100 spectrophotometer. After normalizing, the obtained values for control hemolysis growth, the ratio between the total hemoglobin concentrations within the two circuits was calculated. For example:

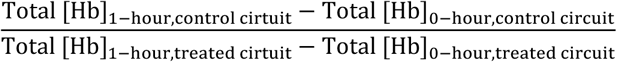

The Cary 100 UV-vis integrating sphere spectrophotometer was employed to determine the blood oxygenation levels by scanning the samples within the 600-900 nm wavelength and fitting them to standard spectra.

### Survival and Failure

Failure of the circuit was defined as the time when the circuit pressure before the oxygenator reaches 200 mmHg which also correlates with very low blood pressure after the oxygenator (less than 10 mmHg). The last collected blood samples were 5 mL/circuit and aliquoted similar to the first 5 mL draw. Survival was defined as lasting six hours without failure. Both reservoirs, oxygenators, tubing, and solutions were replaced and renewed for each experiment.

### Nitrite Dosing Measurement by tri-iodide-based gas phase chemiluminescence

Ozone-based chemiluminescence is a commonly used method capable of detecting up to nanomolar quantities of nitrogen oxides species in solution.^52^ Among various reaction mixtures distinguishing which species are being measured (nitrite, nitrate, or S-nitrosothiols), daily made fresh acidic tri-iodide (I_3_^−^) reagent is a widely used reliable assay to detect nitrite levels.^52,53^ Supernatant samples from the treated system were collected after the nitrite dosing and at the end of the hour before the next infusion.

Three repeats of 5 μL volume of each sample were injected into the purge vessel of the Zysense (formerly Sievers) Nitric Oxide Analyzer (NOA) 280i containing 9 mL tri-iodide reagent at 37°C. The tri-iodide reagent converts the nitrite in a sample into Nitric Oxide which in turn generates a voltage signal inside the NOA. The voltage signal is recorded as a function of time by the software ‘Liquid’ which also measures the area under the voltage curve and converts the area into corresponding nitrite concentration by fitting the area to a nitrite standard curve. The average from the three injections of a sample gives the nitrite concentration for the sample.

### Asynchronous Circuit Testing

Prior to our use of a mixture of packed red blood cells and platelet-rich plasma, we used whole blood in our early testing systems and the systems run sequentially instead of simultaneously. We then switched to the red cell/platelet mixtures since the whole blood had been stored on ice which greatly diminishes platelet function. We include preliminary data using whole blood in this report as well. Before the six simultaneous experiments conducted with the control and treatment circuits, other experiments were conducted as we refined our testing strategy. Four experiments were done with a mixture of packed RBCs with platelet-rich plasma where the untreated and treated circuits were not conducted simultaneously. In two experiments, testing was performed on the control on the first day and on the treatment on the following day. The other two trials were done on the same day but not simultaneously. They started with the control system and after its failure, the treated circuit was tested. In all experiments, calcium or ADP infusions were identical for the control and treatments.

## Results

Several outcomes were documented throughout this study including the changes in pressure as a result of clot formation in the systems, alterations in the number of platelets at the end of the circulation time, blood oxygenation levels and hemolysis changes as a function of time, and percentage improvement in overall circuit survival time in both untreated control and treated systems in all trials. The survival time is defined as the time period in which the system functions under the mean pressure of 200 mmHg during the course of the experiment.

### Circuit Survival

Figure 2 shows examples of the measured pressure before the oxygenator from the control (in black) and the treated (in red) systems on simultaneously running parallel circuits. The treated system involved 100 μM nitrite hourly infusion and red light illumination. The same volume of saline was infused into the control system at the same time as nitrite dosing and mean blood pressure in both systems was recorded every thirty minutes. Panels A-C of Figure 2 show the failure of the untreated circuits (black line shooting vertically upward). Each panel represents platelets and red cells from different donors for experiments on different days and there is variability in circuit survival due to inherent differences in platelet function and other factors.

**Figure 2.**
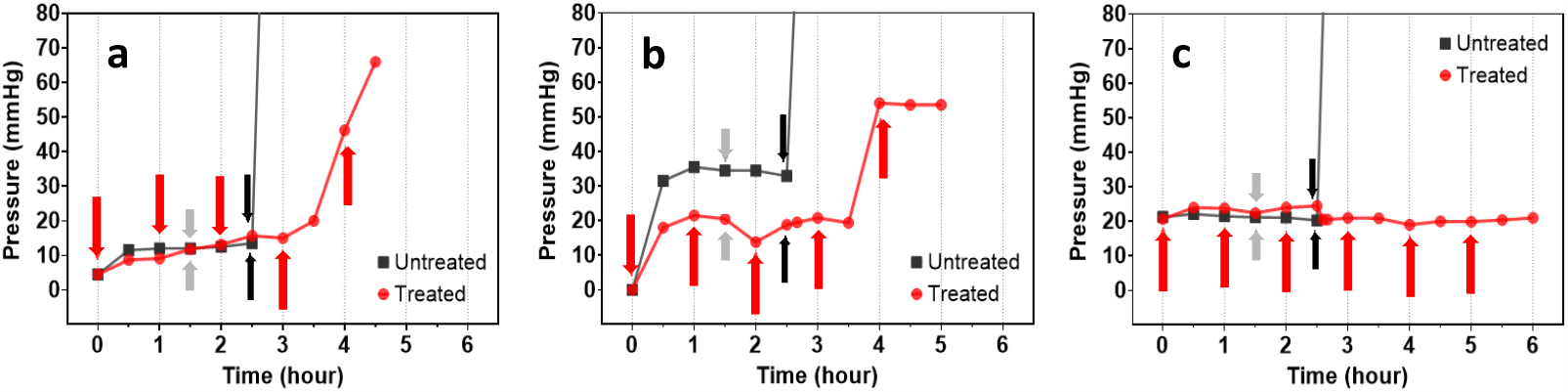
Examples of mean pressure of the untreated control (black) and treated (red) circuits. Consistently, treated circuits survived longer under the same addition of platelet activators as in the untreated circuits. The red arrows are hourly 100 μM nitrite infusion. Gray arrows are for 0.5 μM ADP addition and black arrows are for calcium infusion (2.5 mM in **a** and **b**, 1.5 mM in **c**). The ratio of RBC:saline:PRP is 1.7:1.7:1.

Examined outcomes from six simultaneously paired experiments demonstrate the potential of our dual nitrite and red light treatment as an effective therapy in prolonging circuit survival. In half of the cases, our treatment completely prevented the circuit failure as the final pressure values in the treated systems were maintained in the lower range while the untreated were clotted beforehand. All panels of Figure 3 represent the increased circuit survival time in the treated systems. Figure 3a summarizes the six simultaneously-run experiments. Panel b shows the results of the ten trials using red blood cell mixture with platelets and the outcomes were significant in increasing the circuit survival. These results are a combination of simultaneous and sequential runs including same-day and two-days experiments. Panel c of Figure 3 has all successful experiments combined together. Similar effects of system survival prolongation were seen regardless of the use of whole blood or blood mixture of RBCs plus platelet-rich plasma.

**Figure 3.**
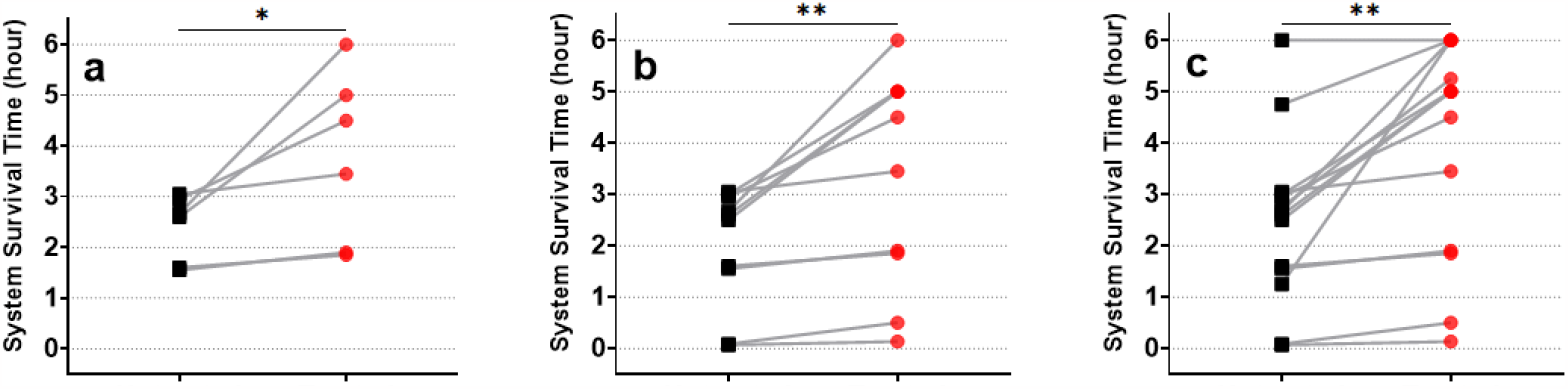
Symbol and line graph of survival proportions. Black squares are untreated controls, and red circles are the treated circuits. Panel **(a)** includes the six paired untreated vs treated tests with RBC+PRP mixture, running at the same time with longer survival time for the circuit treated with nitrite and light illumination (*p < 0.05, n = 6). Panel **(b)** is the combined results of ten experiments using a mixture of red blood cells and platelet-rich plasma (**p = 0.006, n = 10). Panel **(c)** represents all trials from this study, including using whole blood added to the mixture of red cells and platelets (**p > 0.001, n = 14).

Figure 4 is the Kaplan Meier survival curves of the simultaneous tests (panel a, n = 6) and ten combined runs with red blood cells mixed with platelet-rich plasma (panel b). Consistent improvement caused by the dual treatment of nitrite and red light is notable in all panels regardless of the timing of the running systems. Figure 3c represents the survival time enhancement in a total of fourteen gathered trials with whole blood and blood mixture.

**Figure 4.**
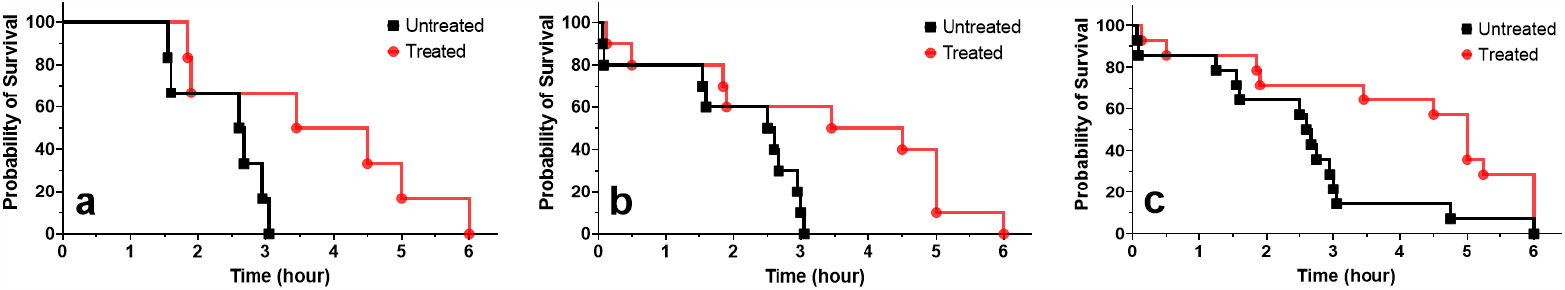
Kaplan Meier survival graph. Panel **a** is dedicated only to the six simultaneous runs. Log-rank (mantel-cox) test reveals a significant difference in the curves with *p = 0.04, n = 6. Panel **b** includes combined results from ten paired experiments using a mixture of packed red cells, saline, and platelet-rich plasma (*p = 0.02, n = 10). Panel **c** is the survival in all fourteen trials regardless of the mixture or the whole blood used in the system (*p = 0.02, n = 14). In all panels, the black lines and symbols are representatives of the untreated control circuit while the red lines and symbols stand for the circuit benefiting from the nitrite/red-light therapy.

Regardless of the survival period, the blood mixture used, and the length of the procedures in each trial, the effects of the combined treatment of nitrite infusing and red light illumination improved circuit survival time in the treated systems. Figure 5 shows the relative improvement in circuit survival. The range was from a minimum of hundred percent to a maximum of six hundred percent compared to the untreated control systems.

**Figure 5.**
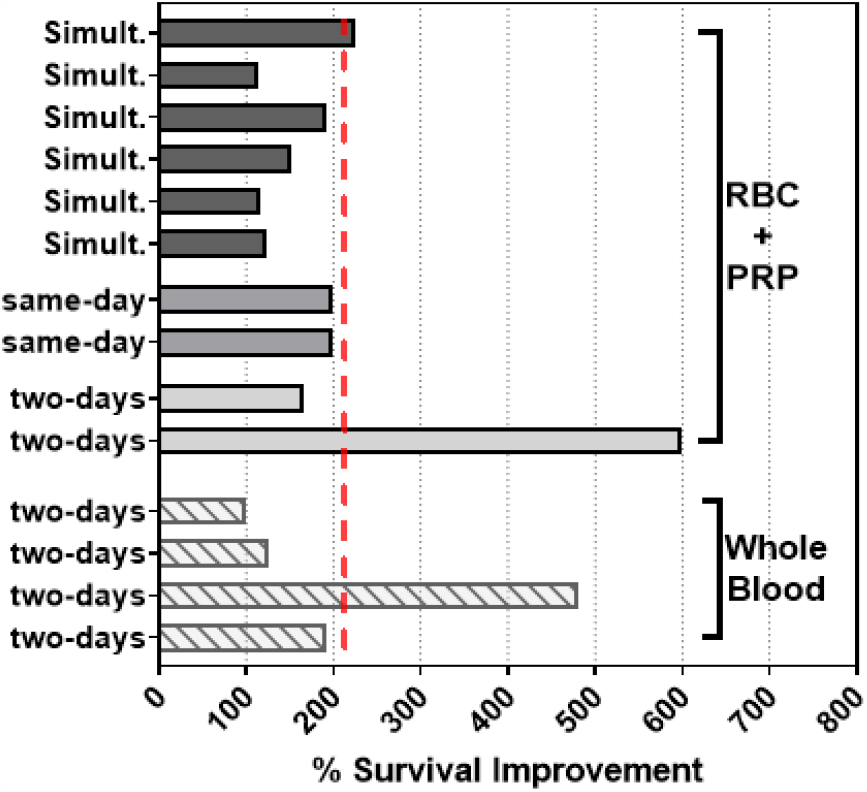
The percentage range of the improvement by the dual nitrite/light treatment. On average, the survival time of the treated system is 213% longer than the control (red dashed line). The bottom four columns are experiments with whole blood while the rest of the data show results of trials done by using a mixture of red blood cells (RBCs) and platelet-rich plasma (PRP).

### Enhanced Platelet Count

In addition to circuit survival, Figure 6a shows the noticeable effects of the treatment on the platelet counts. Platelets become activated during the time course of the procedure, and as a result, the detected number of platelets showed in a Complete Blood Count (CBC) test decreases when blood samples from later time points were compared with the initial platelet count. Averaged data from four CBC tests using the mixture of red cells and platelet-rich plasma showed a 71% decrease in the number of functional platelets in the untreated system collected from the first time point sample (“0 hours”) to the last sample drawn from the circuits (“last hour”) regardless of the length of the circuit survival time. This value for the treated system with nitrite infusion combined with light therapy is 45% loss averaged from four coupled experiments.

**Figure 6.**
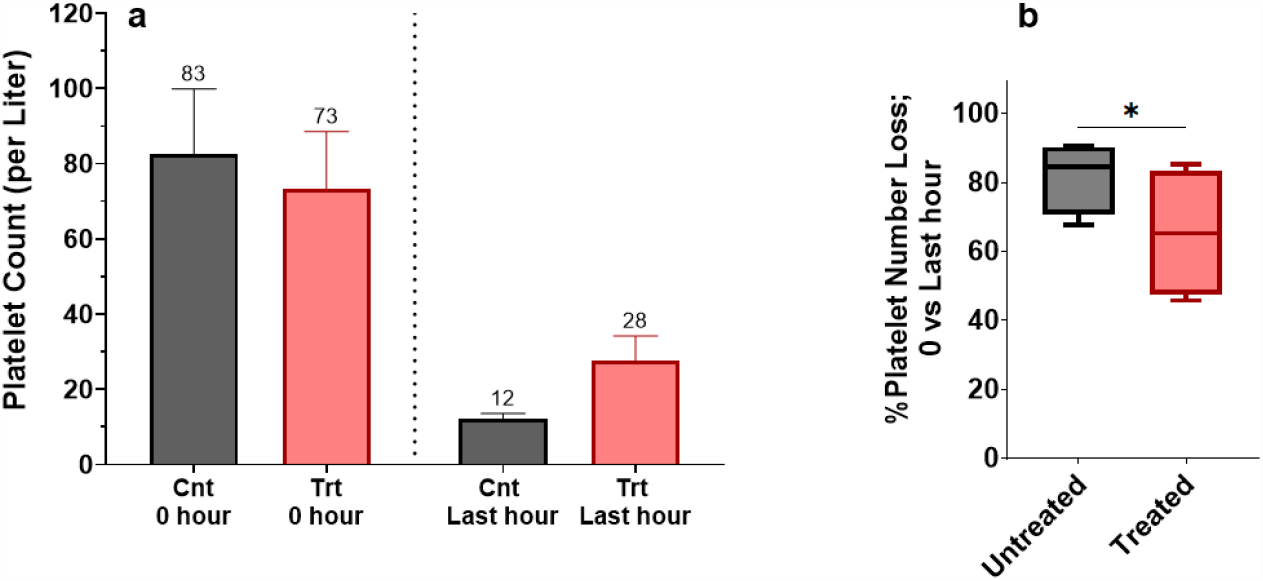
Averaged platelet count and percentage of platelet loss from CBC analysis. Panel **(a)**: Averaged platelet counts from four experiments using RBCs+PRP. The untreated system is the control system (“Cnt”). In the absence of the dual treatment, the untreated systems lost 71% of the platelets between the start to the end of the circuit failure. Treated circuits on the other hand showed only a 45% decrease in the platelet count (n = 4, mixture of packed red blood cells and plasma) proving the positive effects of the nitrite and light therapy on saving functional platelets over time. Panel **(b)**: Percentage of platelet loss at each trial calculated by dividing the change in the number of platelets in the treated circuits by the change in the platelets number in the control systems between zero to last hour of all four experiments. Nitrite combined with red-light treatment significantly reduced the platelet loss shown in percentage format (*p < 0.05, n =4). Untreated: %82 averaged platelet loss; Treated circuit: %65 averaged loss.

Panel b of Figure 6 includes the normalized data in each of the four trials by calculation of the averaged ratio between the final platelet loss in the treated circuit versus the untreated control system.

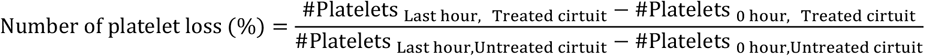

On average, a significant reduction effect in platelet loss was recorded due to the combination therapy of nitrite infusion and red-light treatment.

### Hemoglobin Oxygen Saturation and Improved Hemolysis

Absorption spectra analysis used to determine blood oxygenation levels are shown in Figure 7. Panel A includes the basis absorption spectra of fully oxygenated and 100% deoxygenated hemoglobin samples. Figure 7B is an example of fitting a sample curve to the theory spectrum where most of the sample is at a maximum deoxygenated state. However, this condition is lower than venous oxygen saturation *in vivo*, a hypoxic environment is favorable for enhanced NO bioactivation compared to normoxia.

**Figure 7.**
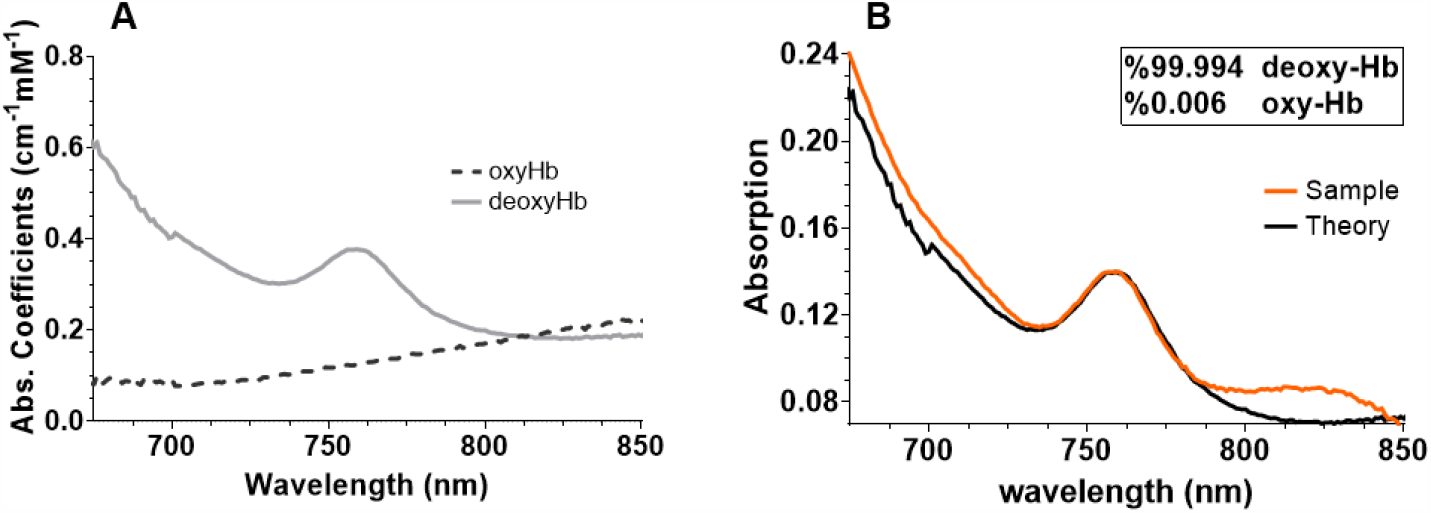
Hemoglobin Oxygen Saturation curves. (**A**) Basis spectra for standard oxygenated Hb (dashed line) and deoxygenated Hb (solid line) within 675-850 nm. (**B**) Sample absorption fitting to the theory spectrum. The sample contains 99% deoxygenated hemoglobin at a concentration of 355 micromolar.

Another improved factor in this study was the degree of hemolysis as the red blood cells rupture over time and circulation. Hemolysis contributes to platelet activation and its intravascular levels have been associated with increased morbidity and mortality.^54^ Hemolysis leads to hemoglobin directly scavenging nitric oxide and releasing ADP which activates platelets.^55,56^ Figure 8 shows the effects of nitrite and red-light therapy on reducing hemolysis levels over the course of six experiments using the mixture of red cells and platelets. On average, a reduction of 2.3 fold was achieved for the treated circuits. Figure 8a shows representative absorption spectra of hemolysis over time in the control system (solid lines) in comparison with the treated circuit (dashed lines). Figures 8 b and c are summarized changes in absorption spectra of blood samples collected hourly until the failure of the circuits.

**Figure 8.**
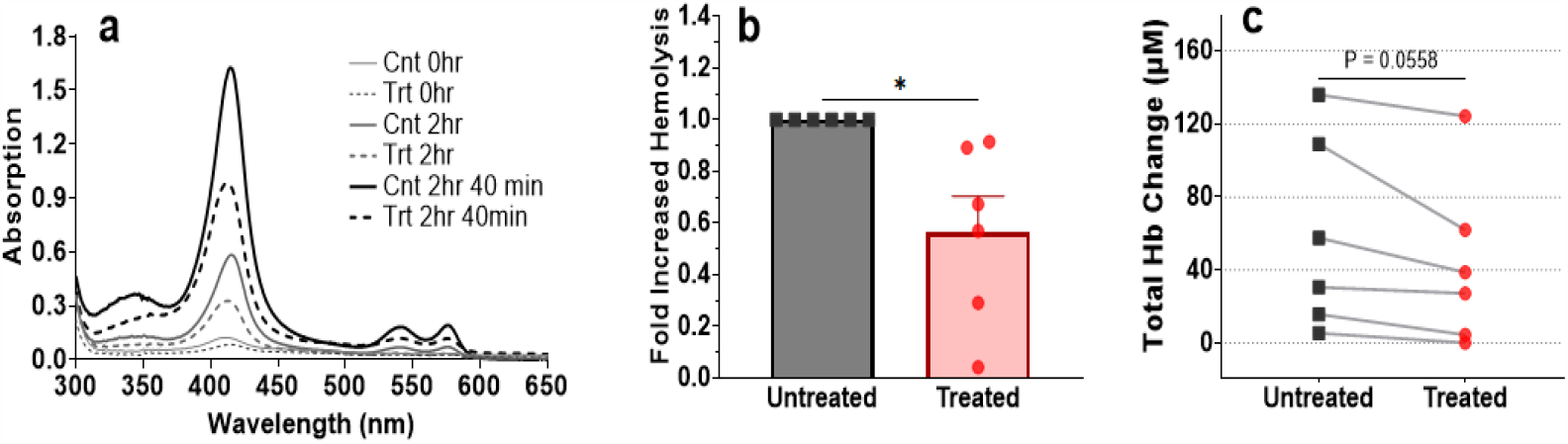
Hemolysis Changes. **(a)** An example of the absorption spectra in the wavelength range of 300-650 nm. The solid lines show the increasing trend of hemolysis over time from the bottom to the top in the untreated control system (“Cnt” labeled) from the zero point to the last sample drawn from that system before complete failure. The dashed lines are the progressing curves of hemolysis changes in the system using the dual therapy matched with the control time points. At each point, the hemolysis is notably lower in the treated circuit. **(b)** Fold increase in hemolysis levels. The average in the treatment was 0.56 when normalized to the related control samples (n = 6, *p = 0.03, a mixture of RBC+PRP). **(c)** Changes in total hemoglobin concentration in six runs with the mixture of blood and platelets (p = 0.056, n = 6).

### Nitrite Dosing Measures

Our nitrite infusions consisted of 0.05 millimoles of nitrite in 500 mL total volume circulating in the circuit. Based on our selected dosage and considering 5 liters of blood in an adult human, the equivalent nitrite dosing infused in a patient would be 0.5 millimoles per hour. Not all the nitrite administered would reach the patient as it is reacted with red blood cells. We measured an average of 80 μM nitrite present in the system after the first circulation (3 minutes immediately after 100 μM hourly nitrite infusion). After one hour of circulation, we measured 28 μM nitrite. The 52 μM averaged reduction occurred hourly as the system was running during the course of the experiments. Figure 9 shows the results from samples of treated circuits.

**Figure 9.**
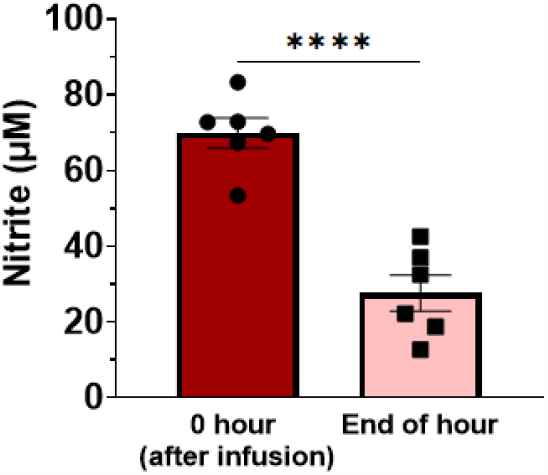
Nitrite level drop in an hour. A chemiluminescence-based nitrite oxide analyzer showed an average of 80 μM nitrite concentrations after each infusion. This value dropped to 28 μM at the end of the hour of blood circulation. These results are collected from the simultaneous trials using RBCs and platelets mixture. (****p < 0.0001, n = 6).

## Discussion and Conclusions

We built a system that consists of two circuits so we can compare control to treatment (or compare two different treatments) run at the exact same time. We have tested the system comparing treatment to control in terms of patency, platelet count, and hemolysis. Our combined treatment of nitrite and light increased circuit survival time, preserved platelet count, and decreased hemolysis. These results are in agreement with earlier versions of our testing strategy where samples were not run simultaneously and when we employed whole blood for testing. On average, the survival period of studied systems was prolonged by 213% in the total of fourteen successful trials. Overall, our experiments show that the combined light/nitrite treatment has potential to prevent device thrombosis.

One limitation of our study is that even with our platelet and red blood cell mixture, we do not exactly reproduce the situation in humans undergoing continuous dialysis. Future experiments should employ freshly drawn whole blood (rather than mixtures of recently drawn red blood cells and platelets), but this requires blood drawing in a clinical setting due to the volumes involved. Another limitation is that although we aimed to achieve a hemoglobin oxygen saturation similar to that for mixed venous blood using nitrogen gas and an oxygenator, we ended up with a lower hemoglobin oxygen saturation. Given that the optimal hemoglobin oxygen saturation for nitrite reduction to NO is around the hemoglobin p50^57^, more work needs to be done to see how large an effect the lower oxygen tension had. The fact that these were only *in vitro*, and not *in vivo* experiments is also a limitation. Future work will include determining the optimal nitrite infusion rates and doses as well as the optimal light illumination intensities. The relative role of the nitrite vs the light needs to be elucidated.

This study focused on potential application to extracorporeal circulation. Possible safety concerns due to the nitrite include hypotension and methemoglobinemia. Pluta et al did a safety study for long-term nitrite infusion and determined that the maximum tolerable dose is 267 μg/kg/hr and dose-limiting toxicity was reached at 446 μg/kg/hr (with toxicity defined as an asymptomatic decrease in blood pressure or increase in methemoglobin).^58^ For a 70 Kg individual, the maximum tolerable dose would be 0.4 mmoles/hr. We were infusing nitrite in our test system at 0.05 mmoles/hr in our *in vitro* circuits. Thus, we could safely infuse at a much higher concentration which could have even bigger effects on preventing local thrombosis. In addition, it is important to note that Pluta et al were infusing directly into volunteers whereas a portion of the nitrite we infuse extracorporeally will be reacted before going into the patients.^57^ There are no known safety issues tied to using far-red light, but safety needs to be assessed in an animal model.

In conclusion, we have shown that the combined application of nitrite and far-red light illumination that we have previously shown to inhibit clotting^25^ has potential in the prevention of device thrombosis.

## Acknowledgments

This work was funded by Wake Forest Innovations and the Translational Science Center. We thank Dr. Anita Saran for helpful discussions.

